# Structural and biochemical characterization of an encapsulin-associated rhodanese from *Acinetobacter baumannii*

**DOI:** 10.1101/2024.02.19.581022

**Authors:** Robert Benisch, Tobias W. Giessen

## Abstract

Rhodanese-like domains (RLDs) represent a widespread protein family canonically involved in sulfur transfer reactions between diverse donor and acceptor molecules. RLDs mediate these transsulfuration reactions via a transient persulfide intermediate, created by modifying a conserved cysteine residue in their active sites. RLDs are involved in various aspects of sulfur metabolism, including sulfide oxidation in mitochondria, iron-sulfur cluster biogenesis, and thio-cofactor biosynthesis. However, due to the inherent complexity of sulfur metabolism caused by the intrinsically high nucleophilicity and redox sensitivity of thiol-containing compounds, the physiological functions of many RLDs remain to be explored. Here, we focus on a single domain *Acinetobacter baumannii* RLD (Ab-RLD) associated with a desulfurase encapsulin which is able to store substantial amounts of sulfur inside its protein shell. We determine the 1.6 Å x-ray crystal structure of Ab-RLD, highlighting a homodimeric structure with a number of unusual features. We show through kinetic analysis that Ab-RLD exhibits thiosulfate sulfurtransferase activity with both cyanide and glutathione acceptors. Using native mass spectrometry and *in vitro* assays, we provide evidence that Ab-RLD can stably carry a persulfide and thiosulfate modification and may employ a ternary catalytic mechanism. Our results will inform future studies aimed at investigating the functional link between Ab-RLD and the desulfurase encapsulin.

## Introduction

Rhodaneses and rhodanese-like domains (RLDs) are a ubiquitous class of enzymes, found across all domains of life, involved in sulfur transfer reactions^1^. The rhodanese family (PF00581) exhibits a conserved protein fold characterized by an α-β-repeat in which a central β-sheet is surrounded by α-helices^2^. Rhodaneses can occur as single standalone domains, as well as multi-domain proteins, containing multiple distinct RLDs or a combination of RLDs and other non-rhodanese domains^1^. Regardless of domain architecture, the function of RLDs is to catalyze transsulfuration reactions, transferring sulfur atoms from small molecule or protein donors to small molecule or protein acceptors, via intermittent persulfide intermediates formed on a conserved cysteine within the RLD active site^3,4^. Rhodanese domains have been implicated in a variety of biological functions, such as the sulfide oxidation pathway in mitochondria^5^, molybdenum cofactor (MoCo) biosynthesis^6,7^, iron-sulfur cluster biogenesis^8^, and thiouridine biosynthesis^9,10^. Multiple bacterial and human rhodanese-containing proteins have also been shown to interact with thioredoxins to mediate persulfidation and enable disulfide exchange, including the single domain rhodanese GlpE from *Escherichia coli*^11^, human TSTD1^12^, and RhdA from *Azotobacter vinelandii*^13^.

In a recent study, we identified a putative single domain rhodanese (Ab-RLD) associated with an encapsulin nanocompartment operon found in *Acinetobacter baumannii*^14^. Encapsulin nanocompartments are self-assembling compartmentalization systems widespread in prokaryotes where cargo enzymes are specifically encapsulated inside 18 to 42 nm icosahedral protein shells^15,16–18^. Encapsulins have been shown to be involved in iron storage ^19–21^, peroxide detoxification ^22,23^, and sulfur metabolism^24^. In the *A. baumannii* system, a cysteine desulfurase enzyme was identified as the cargo enzyme and was shown to be able to sequester and store substantial amounts of elemental sulfur inside the encapsulin shell. We proposed that this desulfurase encapsulin may be involved in redox stress resistance and sulfur storage. The stored sulfur may represent a privileged sulfur pool to be remobilized under specific conditions via a so far unknown mechanism. The presence of the Ab-RLD encoding gene in the genomic neighborhood of the cysteine desulfurase and encapsulin genes may suggests a functional link^14^.

In this study, we report the computational, biochemical, and structural analysis of Ab-RLD. We determine the 1.6 Å x-ray crystal structure of Ab-RLD, highlighting a homodimeric structure with a number of unusual features. We show through *in vitro* assays that Ab-RLD exhibits thiosulfate sulfurtransferase activity with both cyanide and thiol-based sulfur acceptors. Using native mass spectrometry, we provide evidence that Ab-RLD can stably carry a persulfide and thiosulfate modification. Based on our data, we propose that Ab-RLD may employ a ternary catalytic mechanism, instead of the canonical double-displacement mechanism proposed for most rhodaneses. Our results will inform future studies aimed at investigating the link between Ab-RLD and its genomic neighbors, including the encapsulated cysteine desulfurase.

## Results

### Bioinformatic analysis of rhodanese-like domains (RLDs)

To investigate the distribution and diversity of RLDs, and specifically encapsulin-associated RLDs, we carried out a large-scale bioinformatic search using the Enzyme Function Initiative-Enzyme Similarity Tool (EFI-EST). Our EFI-EST search yielded a 10,000-member sequence similarity network (SSN) of putative rhodaneses with more than a dozen major clusters at a sequence identity threshold of 29% (Fig. S1*A*). The predicted RLDs belonged to a single protein family (PF00581). Encapsulin-associated RLDs were confined to a single large and heterogeneous cluster. We next focused our analysis on the cluster containing all encapsulin-associated RLDs and applied a more stringent identity threshold of 49%. After removing any fusion our multi domain rhodaneses by restricting sequence length to 200 amino acids, a fairly homogeneous encapsulin-associated RLD cluster – containing Ab-RLD – as well as several other large and mostly homogeneous clusters of related RLDs, could be obtained (Fig. 1*A*).

The individual clusters generally contained conserved genome neighborhoods and gene synteny with respect to the RLD-encoding genes (Fig. 1*B*). In the two largest clusters (cluster 1 and 2), RLD genes were found next to glutamine synthetase and *O*-succinylhomoserine sulfhydrylase (OSHase) genes, respectively. While glutamine synthetase is not directly involved in sulfur metabolism, hydrogen sulfide (H_2_S) has been implicated in the stimulation of lipid biosynthesis from glutamine^25^. OSHase catalyzes the pyridoxal phosphate (PLP)-dependent conversion of *O*-succinylhomoserine and hydrogen sulfide (H_2_S/HS^-^) to homocysteine. OSHase-associated RLDs are also found in a number of the other clusters, including a rhodanese from *Mycobacterium tuberculosis*, structurally characterized as part of a structural genomics project.

**Figure 1.**
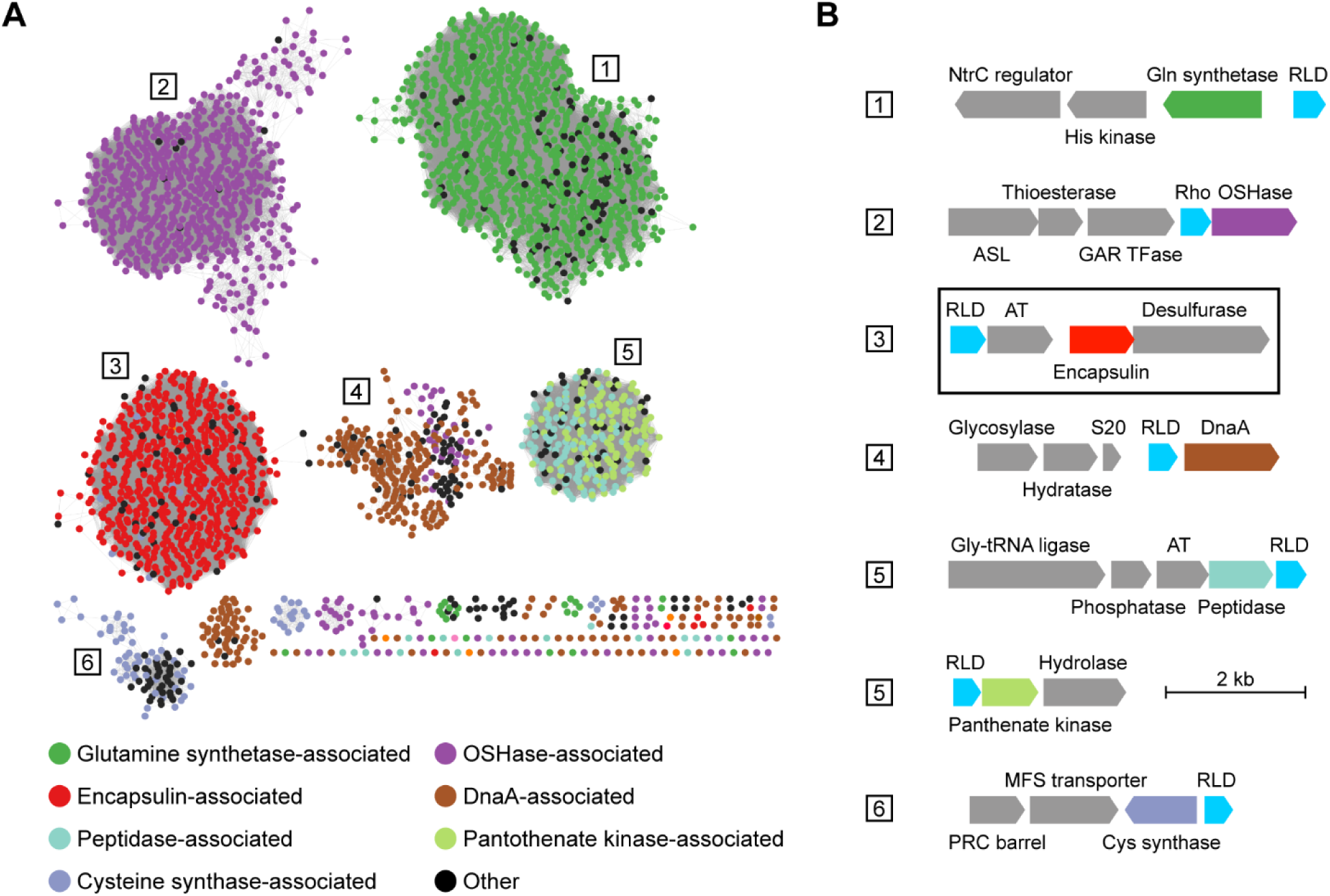
Computational analysis of bacterial RLDs. *A*, sequence similarity network (SSN) of bacterial RLDs with similarity to Ab-RLD and other encapsulin-associated RLDs. Sequence identity cut-off: 49%. Nodes are colored by conserved genome neighborhood features surrounding RLDs. OSHase: *O*-succinylhomoserine sulfhydrylase, DnaA: chromosomal replication initiator protein DnaA. *B*, representative conserved genome neighborhoods found within the six largest SSN clusters. Cluster 3 contains all encapsulin-associated RLDs including Ab-RLD. The Ab-RLD genome neighborhood is highlighted. RLD: rhodanese-like domain, ASL: adenylosuccinate synthetase, GAR TFase: glycinamide ribonucleotide transformylase, AT: acetyltransferase, S20: small ribosomal subunit protein bS20. kb: kilobases.

As rhodaneses have been shown to directly produce H_2_S from thiosulfate with a reducing agent present^26,27^, the role of OSHase-associated RLDs may be to generate H_2_S as a direct substrate for OSHases. Cluster 3 contains all encapsulin-associated RLDs, generally found in conserved four-gene operons (Fig. 1*B*). Besides the RLD and encapsulin genes, these operons generally encode a serine-*O*-acetyltransferase (AT) and a cysteine desulfurase (CD). While CD functions as an encapsulin cargo protein and allows the sequestration of elemental sulfur inside the encapsulin protein shell^14^, the function of AT is currently unknown. In general, ATs catalyze the production of *O*-acetylserine from acetyl-CoA and serine. *O*-acetylserine is the direct precursor for cysteine biosynthesis via cysteine synthase^28^. Both ATs and encapsulin-associated RLDs may be involved in the mobilization of sulfur stored inside the encapsulin shell. The genome neighborhoods of RLDs found in clusters 4 and 5 are more varied and no obvious connection to sulfur metabolism can be proposed. RLDs in cluster 6 on the other hand are encoded next to cysteine synthase genes. These RLDs could again be involved in generating H_2_S as a substrate for cysteine synthases for the transformation of *O*-acetylserine to cysteine^28^.

Using sequence alignments, we next focused on comparing Ab-RLD with other RLDs contained within our dataset. As sulfurtransferases, rhodaneses are able to accept a sulfur atom from a donor, store it intermittently as a persulfide intermediate on an active site cysteine, before passing it on to a nucleophilic acceptor^29^. Sequence logos of the active site residues of RLDs from different SSN clusters highlight the strictly conserved active site cysteine (Fig. S1*B*). All active sites exhibit a general CRSG/AxR motif. Notably, the majority of encapsulin-associated RLDs contain a positively charged lysine at position 5 whereas RLDs from other clusters often exhibit small or hydrophobic residues at this position, although in Ab-RLD an asparagine residue is found at position 5. We further observed that Ab-RLD and most encapsulin-associated RLDs contain an extended unannotated N-terminus, not present in most other RLDs such as the structurally characterized OSHase-associated RLD from *M. tuberculosis* (Fig. S1*C*). This is also reflected in the average length of encapsulin-associated RLDs which, at 162 residues, is larger than the average lengths of RLDs from other clusters that range from 125 to 153 residues (Fig. S1*D*).

### Structural characterization of Ab-RLD highlights conserved and divergent features

The x-ray crystal structure of Ab-RLD was obtained at 1.6 Å resolution (Fig. 2*A*, Table S1). The final model contained two chains in the asymmetric unit (ASU). Size exclusion chromatography further confirmed that Ab-RLD exists as a homodimer in solution (Fig. S2*A*-*C*). The Ab-RLD monomer exhibits an α-β-repeat fold typical of PF00581 rhodanese domains (Fig. 2*B*). In the context of the Ab-RLD homodimer, the two active sites are located at opposite sides of the protein complex (Fig. 2*A* and *B*). The first 10 amino acids of the N-terminal extension found in Ab-RLD are visible in only one chain of the ASU (Fig. 2*A*). An N-terminal α helix (α1), usually not found in PF00581 rhodanese domains, is clearly resolved in both chains (Fig. 2*A*).

**Figure 2.**
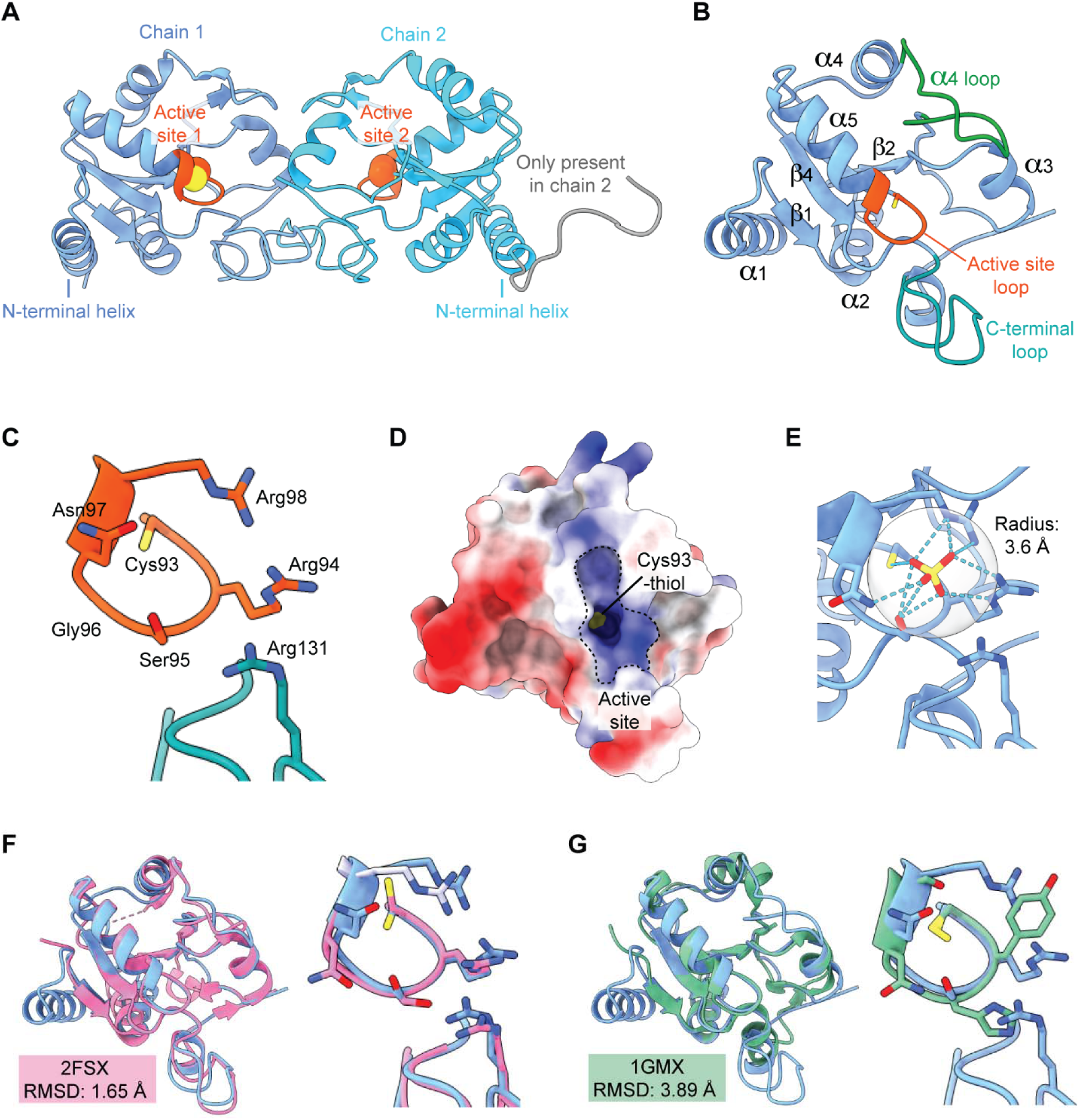
Structural analysis of Ab-RLD. *A*, homodimer of Ab-RLD shown in ribbon representation. The two active sites are highlighted in orange – including the active site cysteine (Cys93), shown as spheres. The N-terminal helices, found mostly in encapsulin-associated RLDs, are highlighted. Ten residues, only resolved in chain 1, are shown in gray. *B*, a single Ab-RLD monomer in ribbon representation is shown with secondary structural elements labeled. The active site is shown in orange while two features not found in all RLDs – the a4 loop and the C-terminal loop – are highlighted in green and cyan, respectively. *C*, detailed view of the extended active site, including the active site loop and the catalytic Cys93. Residues proposed to be important for Ab-RLD function are shown as stick models. *D*, electrostatic surface representation of Ab-RLD monomer in the same orientation as in *B*. The strongly positively charged active site and the thiol of Cys93 are highlighted. *E*, active site with modeled sulfate ion coordinated by active site loop residues. Hydrogen bonds within 3.6 Å (transparent sphere) of the sulfate ion are shown in cyan. *F*, left: Structural alignment of monomers of *M. tuberculosis* RLD (PDB ID: 2FSX, pink) and Ab-RLD (blue). Right: Active site comparison. *G*, left: Structural alignment of monomers of *E. coli* GlpE (PDB ID: 1GMX, green) and Ab-RLD (blue). Right: Active site comparison.

The Ab-RLD active site is located at the loop connecting strand β3 and helix α5 which contains the active site cysteine (Cys93) (Fig. 2*C*). In addition to Cys93, residues Arg94, Ser95, Gly96, Asn97, and Arg98, constitute the loop between β3 and α5 and form a positively charged surface-exposed pocket that may allow Ab-RLD to interact with a variety of different sulfur donors and acceptors. (Fig. 2*D*). In addition, Arg131, not part of the active site loop, seems to be part of the active site, contributing to the overall strong positive charge surrounding Cys93 (Fig. 2*C*). An additional density with tetrahedral geometry was observed inside the active site pocket, coordinated by multiple active site loop residues (Fig. 2*E* and Fig. S2*D*). The density was modeled as a sulfate ion, as the crystallization conditions contained sulfate.

A number of bacterial RLDs have been structurally characterized within the context of different structural genomics projects, however, little is known about their biochemical or physiological function. Comparing Ab-RLD to other structurally characterized RLDs of high sequence similarity, we found that the abovementioned *M. tuberculosis* RLD (PDB ID: 2FSX, sequence identity: 44%) exhibits high overall structural similarity with an RMSD of 1.65 Å and an identical active site composition, including the active site loop and the equivalent of Arg131 in Ab-RLD (Fig. 2*F*). A clear difference between the two proteins is the N-terminal α1 helix of Ab-RLD, which is not present in the mycobacterial protein. The biochemically and structurally characterized rhodanese GlpE from *Escherichia coli* (PDB ID: 1GMX) – whose biological function is still unknown – exhibits a sequence identity of 24% while a structural alignment with Ab-RLD yields an RMSD of 3.89 Å (Fig. 2*G*)^30^. The N-terminal α1 helix is not present in GlpE. In contrast to the mycobacterial RLD and Ab-RLD, GlpE lacks the extended loop after helix α4 (α4 loop) and the C-terminal loop (Fig. 2*B*). When present, the loop extending from α4 lies above the active site loop and creates an extended active site pocket (Fig. 2*D*). The active site loop found in GlpE is substantially different from Ab-RLD with only two residues being conserved corresponding to Cys93 and Asn97 in Ab-RLD (Fig. 2*G*). Based on previous reports, active site loop residues can likely influence substrate specificity, suggesting that GlpE and Ab-RLD might utilize different sulfur donors or acceptors^30^.

### Ab-RLD is a thiosulfate sulfurtransferase

We next sought to investigate the sulfurtransferase activity of Ab-RLD through *in vitro* assays. The canonical *in vitro* approach to confirm the sulfurtransferase activity of rhodaneses is the thiosulfate:cyanide sulfur transfer assay^4^. Briefly, thiosulfate is used as an efficient sulfide donor to rhodaneses, and after binding to the active site, transfers sulfide to the active site cysteine of the rhodanese in question producing a persulfide intermediate. Next, the sulfide is transferred to cyanide, acting as a nucleophilic small molecule acceptor, to produce thiocyanate. Thiocyanate levels can then be determined colorimetrically by adding ferric iron to produce a ferric thiocyanate complex with a strong and characteristic absorption at 460 nm^31^. It is important to note that this assay is useful for evaluating general sulfur transfer activity, but may not necessarily bear physiological significance, as thiosulfate is a relatively minor sulfur species in the intracellular environment^3^. Similarly, cyanide is an excellent sulfur acceptor, but it has been argued that the primary function of most rhodaneses is unlikely to be the detoxification of cyanide^3^.

**Figure 3.**
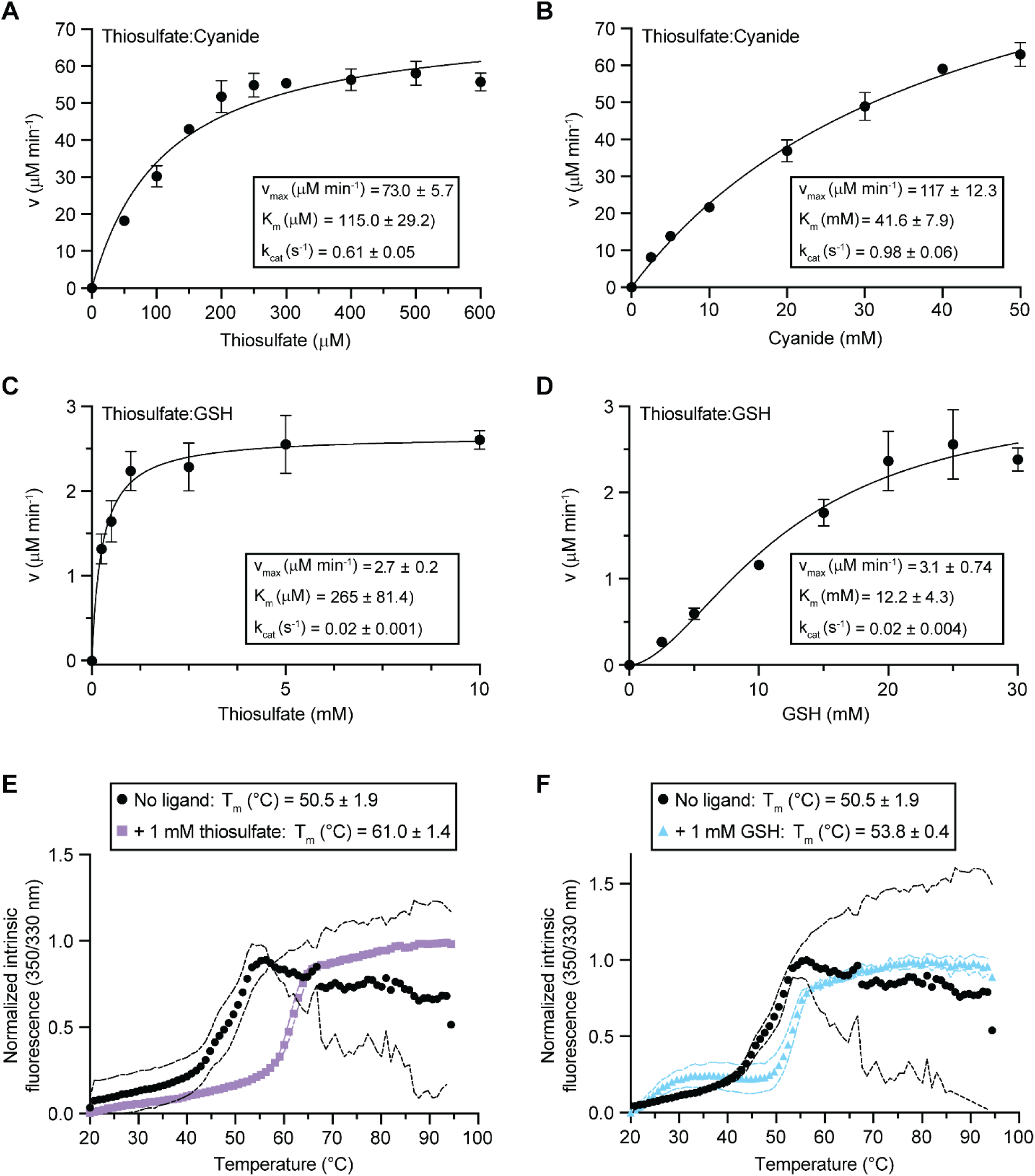
Ab-RLD kinetics and stability. *A*, Thiosulfate:Cyanide sulfur transfer kinetics of Ab-RLD using 50 mM constant cyanide and varying concentrations of thiosulfate. Kinetic parameters are shown. v_max_: maximal velocity, K_m_: Michaelis constant, k_cat_: turnover number. Error bars represent standard deviations of three independent reactions. *B*, Thiosulfate:Cyanide sulfur transfer kinetics of Ab-RLD using 5 mM constant thiosulfate and varying concentrations of cyanide. *C*, Thiosulfate:GSH sulfur transfer kinetics of Ab-RLD using 40 mM constant GSH and varying concentrations of thiosulfate. GSH: glutathione. *D*, Thiosulfate:GSH sulfur transfer kinetics of Ab-RLD using 5 mM constant thiosulfate and varying concentrations of GSH. *E*, melting curves of Ab-RLD in the presence (purple) and absence (black) of thiosulfate. Derived melting temperatures (T_m_) are shown. Curve envelops shown as dotted lines represent standard deviations of triplicate experiments. *F*, melting curves of Ab-RLD in the presence (cyan) and absence (black) of GSH. Derived melting temperatures (T_m_) are shown.

Activity assays using the thiosulfate and cyanide donor-acceptor pair show that Ab-RLD can transfer sulfur from thiosulfate to cyanide and can thus be classified as a sulfurtransferase (Fig. 3*A* and *B*). We obtained k_cat_ values of 0.608 s^-1^ and 0.975 s^-1^ when varying thiosulfate and cyanide, respectively. The corresponding Michaelis constants (K_m_) are 114.9 μM and 41.6 mM for thiosulfate and cyanide, respectively. The kinetic parameters obtained for Ab-RLD are within the same order of magnitude as reported for other single domain rhodaneses which can vary substantially^32^. We carried out additional *in vitro* assays with an alternative sulfur acceptor, namely, glutathione (GSH). GSH is the primary small molecule reducing agent in most cells^33^ and a potentially physiologically relevant small molecule sulfur acceptor for RLD-mediated sulfur transfer reactions^32^. Upon sulfur transfer, GSH is converted to glutathione persulfide (GSSH). GSSH has been suggested to act as a redox regulator ^34,35^ and readily reacts with thiols, including GSH, to produce a disulfide, for example glutathione disulfide (GSSG), with the concomitant evolution of H_2_S^31^. In our *in vitro* assays, H_2_S evolution was measured via the addition of divalent lead ions (Pb^2+^) resulting in insoluble lead sulfide (PbS) precipitant which can be detected turbidimetrically at 390 nm^31^. *In vitro* Ab-RLD sulfur transfer assays using thiosulfate as the sulfur donor showed that GSH is a competent sulfur acceptor (Fig. 3*C* and *D*). We observed k_cat_ values of 0.015 s^-1^ and 0.017 s^-1^ when varying thiosulfate and GSH, respectively. These values are ∼10-fold lower than the previously observed thiosulfate:cyanide transfer activity. Similar to the thiosulfate:cyanide donor-acceptor pair, the K_m_ value for thiosulfate is substantially lower than that of the acceptor GSH – 265 μM and 12.2 mM, respectively. Although the k_cat_ values observed when using GSH as an acceptor were lower than the ones measured when using cyanide, the K_m_ value of GSH was ∼3.5-fold lower than that of cyanide, potentially suggesting that GSH is a likelier candidate for a physiologically relevant sulfur acceptor than cyanide for Ab-RLD. Our GSH acceptor assays show that Ab-RLD is able to produce H_2_S which has also been reported for other RLDs^27, 32^. H_2_S may be used as a substrate in H_2_S/HS^-^-dependent reactions – as described above for OSHase and cysteine synthase – or otherwise function as a signaling molecule^3,36^. In the context of the encapsulin desulfurase system, H_2_S produced by Ab-RLD could be involved in the re-mobilization of stored sulfur via an as of yet unknown mechanism.

We further investigated the binding of the sulfur donor thiosulfate and the sulfur acceptor GSH to Ab-RLD using thermal ramps and internal fluorescence measurements. In the presence of 1 mM thiosulfate, we observed a shift in melting temperature of ΔT_m_ = +10.5°C (Fig. 3*E*) while incubation of Ab-RLD with 1 mM GSH resulted in a ΔT_m_ of +3.3°C (Fig. 2*F*). Based on our activity and structural data, this disparity between thiosulfate and GSH binding to Ab-RLD may be expected. The sulfite moiety of the thiosulfate ion could favorably interact with the arginine residues in the active site loop (Arg94 and Arg98) and Arg131 resulting in the stabilization of the Ab-RLD-thiosulfate complex. To investigate the specificity of the observed increase in T_m_, we carried out additional thermal ramp experiments in the presence of sulfate and phosphate, negatively charged ions with a similar structure compared to thiosulfate. We observed a small thermal shift of ΔT_m_ = +2.2°C in the presence of 2 mM sulfate and no discernable shift using 2 mM phosphate (Fig. S3). These results imply that thiosulfate binds more strongly to Ab-RLD than other anions of similar geometry and size.

### Mass spectrometric analysis shows that Ab-RLD can carry persulfide and thiosulfate

Sulfurtransferases like RLDs are generally characterized by their ability to stably carry persulfides, such as the human MOCS3-RLD^7^ and *Azotobacter vinelandii* RLD^37^. To investigate if Ab-RLD can likewise stably carry persulfide, we subjected Ab-RLD to native mass spectrometric analysis (Fig. 4*A* and Fig. S4). Initially, Ab-RLD was purified in the presence or absence of reducing agents to explore whether it may by persulfidated during heterologous expression in *E. coli*. For *in vitro* sulfur loading, samples were incubated with a molar excess of thiosulfate as a sulfur donor. For sulfur transfer reactions, both thiosulfate and cyanide were added successively.

The resulting mass spectrum of the as-purified oxidized protein sample showed a majority peak for native Ab-RLD (expected monoisotopic mass = 20,877 Da), as well as additional peaks with mass shifts of +32 Da and +48 Da (Fig. 4*B*). A shift of +32 Da is consistent with either the addition of two oxygens to the protein or a cysteine persulfide (Cys-S-SH) while a shift of +48 Da would be consistent with the addition of three oxygens or a persulfide plus a separate single oxygen atom. To discriminate between these possibilities, we incubated the oxidized protein sample with cyanide, which should not be able to remove oxygens, but will revert a persulfide back to a thiol. In the resulting spectra, we observed mass shifts of +16 Da and +32 Da, but no shift of +48 Da (Fig. 4*C*). The abundance of the +16 Da peak is in good agreement with that of the +48 Da peak observed before the addition of cyanide (Fig. 4*B*). This may indicate that the +48 Da peak consists of a persulfide in addition to a separate single oxygen atom and was quantitatively converted to the +16 Da peak in the presence of cyanide. The abundance of the +32 Da peak decreased as well after cyanide addition, suggesting that part of it represents a persulfidated Ab-RLD. For Ab-RLD samples purified under reducing conditions, we observed a single majority peak consistent with native Ab-RLD (Fig. 4*D*). When thiosulfate was added as a sulfur donor, a relatively complex mass spectrum containing likely sodium adducts (+ 21 Da and + 46 Da), a potential persulfide (+33 Da), as well as a majority peak of +112 Da, could be observed (Fig. 4*E*). This majority peak is consistent with the mass of a thiosulfate adduct (Cys-S-S-SO_3_ ^-^), potentially suggesting that it can be stably bound to the active site cysteine. Addition of cyanide resulted in the complete removal of the +112 Da peak and the +33 Da peak (Fig. 4*F*), consistent with these peaks representing thiosulfate-and persulfide-modified Ab-RLD, respectively. Together, our mass spectrometry data suggests that Ab-RLD can stably carry both persulfide and thiosulfate. The formation of a stable thiosulfate-modified Ab-RLD further suggests that Ab-RLD may not follow a double-displacement mechanism, as generally observed for RLD sulfurtransferases^1^.

**Figure 4.**
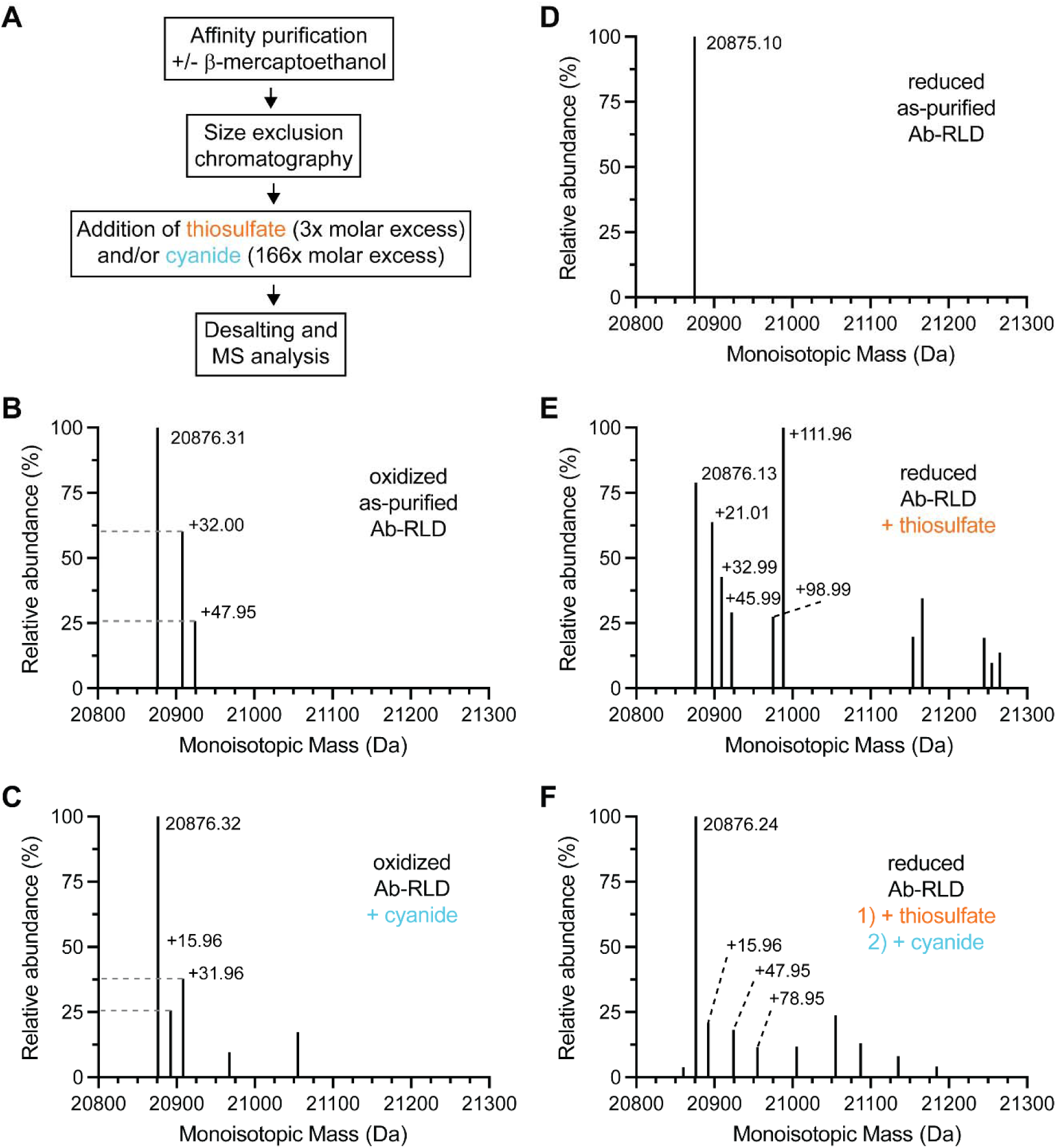
Detection of persulfidated and thiosulfate-modified Ab-RLD via native mass spectrometry. *A*, sample preparation workflow for native mass spectrometric analysis of Ab-RLD. *B*, native mass spectrum of Ab-RLD purified under non-reducing conditions. Relevant peaks are labeled highlighting mass shifts with respect to the native Ab-RLB peak (20,876 Da). *C*, native mass spectrum of Ab-RLD purified under non-reducing conditions with added cyanide. *D*, native mass spectrum of Ab-RLD purified in the presence of the reducing agent b-mercaptoethanol (reducing conditions). *E*, native mass spectrum of Ab-RLD purified under reducing conditions with added thiosulfate. *F*, native mass spectrum of Ab-RLD purified under reducing conditions with successively added thiosulfate and cyanide. Any unannotated mass peaks could not be clearly assigned.

### Ab-RLD may follow a ternary catalytic mechanism

To investigate the sulfur transfer mechanism used by Ab-RLD, we carried out *in vitro* assays monitoring the formation of sulfite from thiosulfate-modified Ab-RLD in single turnover experiments. If Ab-RLD follows a double displacement mechanism, sulfite should be formed from thiosulfate in the presence of Ab-RLD alone (Fig. 5*A*). On the other hand, a ternary mechanism would imply that sulfite formation can only occur in the presence of a sulfur acceptor like cyanide. To detect sulfite in our assays, we used acidified and alkylated *p*-rosaniline which will form a *p*-rosaniline sulfonic acid complex in the presence of sulfite which can be detected colorimetrically at 570 nm (Fig. 5*A*)^27^. We performed single turnover experiments with 50 μM Ab-RLD, from which we would expect to produce up to 50 μM sulfite when supplied with a large excess of thiosulfate sulfur donor. We did not observe any above background sulfite formation in these assays (Fig. 5*B*). Meanwhile, if the sulfur acceptor cyanide was additionally added, substantial sulfite production, indicating multiple turnovers, could be observed, suggesting a ternary instead of a double displacement mechanism for Ab-RLD (Fig. 5*B*).

**Figure 5.**
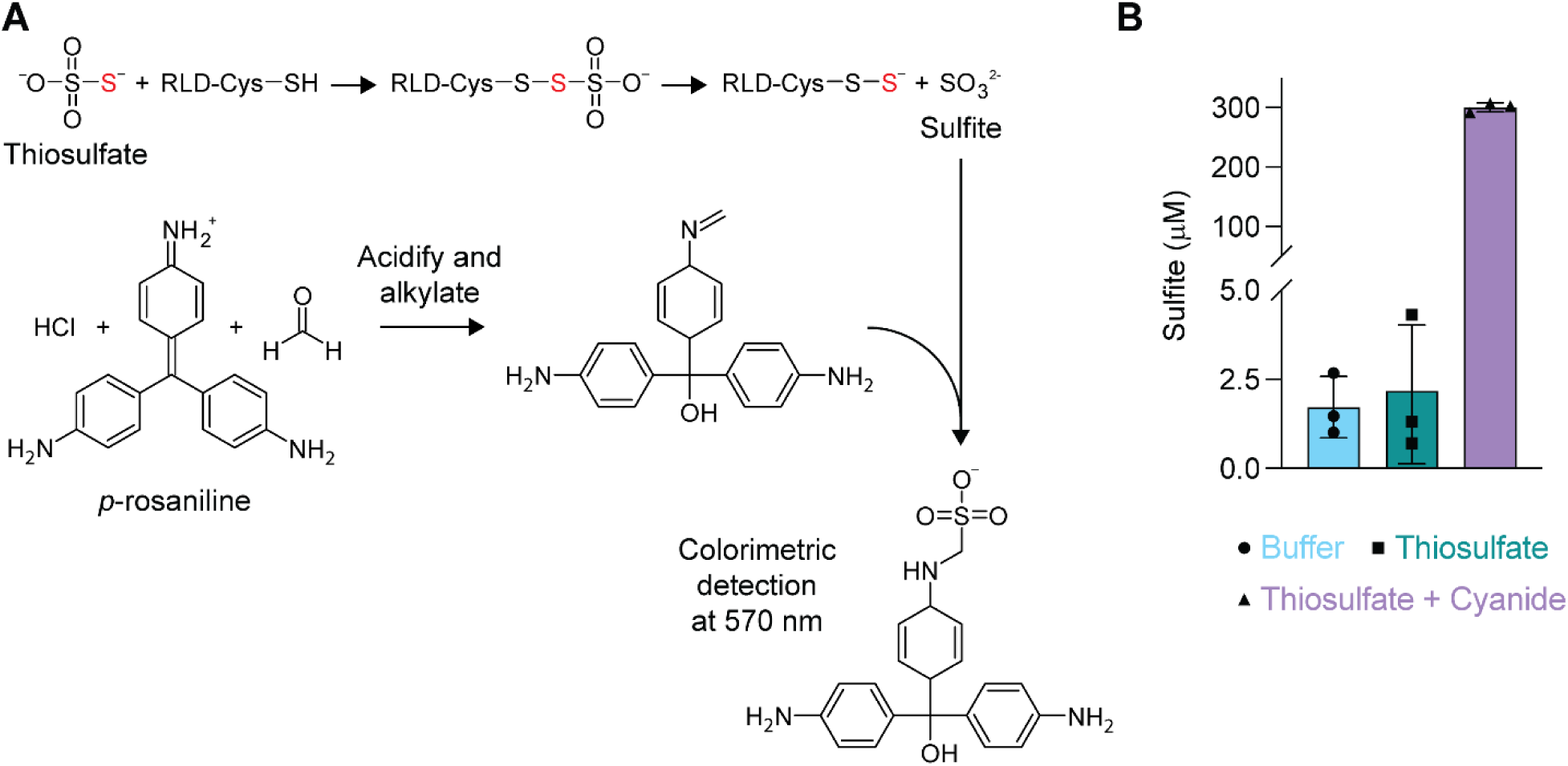
Sulfite production by Ab-RLD in the presence of thiosulfate. *A*, detection scheme for sulfite production by Ab-RLD. *B*, sulfite production by Ab-RLD in the presence of thiosulfate (green) or thiosulfate and cyanide (purple).

## Discussion

In this study, we biochemically and structurally characterize the encapsulin-associated rhodanese-like domain Ab-RLD found in *A. baumannii*, which shares sequence similarity with other single domain RLDs putatively involved in a variety of redox and sulfur metabolism-related metabolic functions^25–28^. Sequence similarity network analysis showed that encapsulin-associated RLDs can be sorted into a homogenous sequence cluster, distinct from other single domain RLDs. While some RLD clusters exhibit conserved RLD genome neighborhoods encoding proteins related to sulfur metabolism, no such connection could be made for other clusters. The gene coding for Ab-RLD is found in a conserved operon encoding three other components, a serine-*O*-acetyltransferase (AT), cysteine desulfurase (CD), and an encapsulin shell protein. It was previously suggested that Ab-RLD may play a role in remobilizing sulfur stored inside the encapsulin protein shell^14^. Encapsulin-associated RLDs, including Ab-RLD, are generally longer than other single domain RLDs, exhibiting an N-terminal unannotated extension not found in other RLDs. As terminal unstructured extensions are known to be important for cargo encapsulation inside encapsulin protein shells, it could be assumed that this N-terminal extension in Ab-RLD might serve a similar function. However, it was recently shown that Ab-RLD does not represent an encapsulin cargo protein^14^. Structural comparisons between Ab-RLD and other structurally characterized single domain RLDs revealed that Ab-RLD exhibits an N-terminal α-helix located in the N-terminal extension, not part of the canonical RLD fold. This structured N-terminal helix may be involved in mediating protein-protein interactions necessary for releasing stored sulfur from within the encapsulin shell. Two other characterized single domain RLDs – Sud from *Wolinella succinogenes*^38^ and AQ1599 from *Aquifex aeolicus*^39^ – contain N-terminal a-helical extensions which have been proposed to be important for translocation to the periplasm.

We further showed that Ab-RLD is capable of sulfur transfer reactions using thiosulfate as a donor, as well as cyanide or glutathione as acceptors. Given the low apparent turnover numbers derived from our kinetic analyses, it is likely that the physiological donor-acceptor pairs of Ab-RLD remain to be discovered. Nonetheless, the substantial thermal shift observed when Ab-RLD is incubated with thiosulfate, but not with sulfate or phosphate, could suggest that thiosulfate or a similar molecule may represent a physiological substrate of Ab-RLD.

Our native mass spectrometry analysis showed that Ab-RLD can stably carry persulfide and thiosulfate. A similar behavior has previously been reported for the *E. coli* RLD PspE which was shown to be able to carry different persulfide (Cys-S-S_n_-SH) and sulfonate (Cys-S-S_n_-SO_3_^-^) modifications, simply as a result of heterologous overexpression^40^. However, in contrast to PspE, only a single persulfide (Cys-S-SH) and sulfonate (Cys-S-S-SO_3_^-^) species could be detected for Ab-RLD. Further, thiosulfate modifications could only be observed after *in vitro* incubation. While PspE has been reported to employ a double displacement mechanism, observed for the majority of RLD sulfutransferases, Ab-RLD may employ a ternary catalytic mechanism as suggested by our single turnover sulfite detection assays. Ternary mechanisms for RLDs have previously been observed for the human thiosulfate:glutathione sulfurtransferase TSTD1 and its yeast ortholog RDL1^41^.

In conclusion, our work lays the foundation for future biochemical, structural, and physiological studies aimed at deciphering how sulfur deposited inside encapsulin shells can be remobilized, a process that potentially involves the now confirmed sulfurtransferase Ab-RLD.

## Experimental procedures

### Bioinformatic analyses of RLDs

The amino acid sequence of Ab-RLD (A0A009HBQ7) was used to initiate SSN generation using EFI-EST^42,43^ with a query e-value of 5 and a maximum number of retrieved sequences of 10,000. The initial 10,000 member SSN (Fig. S1) was generated using Cytoscape v3.10.1^44^. Edges below a sequence identity of 29% were deleted and nodes clustered via the yFiles organic layout algorithm. The 3,187-member cluster containing Ab-RLD was used for further analysis. Multi domain RLDs were removed from the dataset by deleting all nodes with a sequence length >200 residues. The remaining sequences were now clustered again by applying a more stringent sequence identity threshold of 49% to produce the final SSN (Fig. 1*A*). To compute genomic neighborhoods the EFI-GNT server was used. Nodes were colored based on the presence of select conserved RLD neighbors.

Active site sequence logos were created based on Clustal Omega^45^ sequence alignments in Geneious Prime v2023.2.

### Molecular cloning, protein production, and protein purification

The Ab-RLD expression construct was designed to carry an N-terminal His_6_-tag, followed by a Tobacco Etch Virus (TEV) protease cleavage site (ENLYFQG) and a GSG linker. The Ab-RLD construct was codon optimized for *E.coli* K-12 using the Integrated DNA Technologies (IDT) codon optimization tool and synthesized via IDT as a gBlock fragment with appropriate overhangs for insertion into the PacI and NdeI digested pETDuet-1 vector (Table S1 and S2). The final plasmid encoding Ab-RLD was assembled via Gibson Assembly. Subsequently, *E. coli* BL21 (DE3) cells were transformed with the Ab-RLD expression plasmid and the plasmid sequence-verified via Sanger sequencing.

For Ab-RLD protein production, *E. coli* BL21 (DE3) cells carrying the Ab-RLD expression plasmid were grown in TB medium at 37°C and 200 rpm to an OD_600_ of 0.8 and induced using 1 mM IPTG. Induced cells were incubated at 18°C and 200 rpm for 18 h. Cells were harvested by centrifuging at 5,000 *g* for 20 min at 4 °C, flash frozen in liquid nitrogen, and stored at -80°C until further use.

All protein purifications steps were carried out at 4°C unless stated otherwise. Frozen cell pellets were thawed and sonicated in lysis buffer containing 20 mM Tris pH 8.0, 300 mM NaCl, 20 mM imidazole, 3 mM b-mercaptoethanol (omitted for non-reducing purifications of Ab-RLD for downstream mass spectrometric analysis), 0.1 mg/mL lysozyme and 0.1 mg/mL DNAse I, and 1x Pierce protease inhibitor mix. The lysate was then pelleted via centrifugation at 8,000 *g*, and Ni-NTA resin equilibrated in binding buffer (lysis buffer without lysozyme, DNAse I, and protease inhibitor mix) was added to the supernatant and incubated for 2 h under agitation. After incubation, the solution was applied to a Bio-Rad glass column, followed by a washing step with 10 column volumes of binding buffer and 10 column volumes of binding buffer with 20 mM imidazole. The protein was eluted with 250 mM imidazole. To remove the His_6_-tag, 1 mg of tobacco etch virus (TEV) protease was added to 40 mg of Ab-RLD and was subsequently dialyzed against 20 mM Tris pH 8.0, 150 mM NaCl, and 0.2 mM TCEP for 1 h at room temperature, and then at 4°C for 18 h. TEV-cleaved Ab-RLD and TEV protease were separated via Ni-NTA purification as described above. After TEV removal, Ab-RLD was subjected to size exclusion chromatography using a Superdex 200 Increase 10/300 GL in 20 mM Tris pH 8.0 and 150 mM NaCl. Resulting purified Ab-RLD was immediately used for downstream experiments.

### Protein crystallization and structure determination

All crystallization experiments were carried out at the Center for Structural Biology at the Life Sciences Institute at UM. For Ab-RLD crystallization, a buffer exchange into 10 mM HEPES pH 7.5 and 100 mM NaCl was carried out via size exclusion chromatography on a Superdex 200 Increase 10/300 GL. Ab-RLD fractions were then concentrated to 15 mg/mL and kept on ice until crystal tray setup. Trays were set using an Art Robbins Instruments Gryphon microfluidic robot, dispensing 50 μL of crystallization cocktail, and mixing 0.5 μL of cocktail with 0.5 μL of Ab-RLD using the following crystallization screens with sitting drop vapor diffusion: PEG I and II (Nextal Biotechnologies), as well as Hampton Index and JCSG+ (Hampton Research). Crystals used for diffraction experiments were obtained in 0.1 M ammonium sulfate, 0.1 M sodium acetate pH 4.8, and 20% PEG3350, and were harvested by soaking in crystallization cocktail supplemented with 10% ethylene glycol before flash freezing in liquid nitrogen. Diffraction data was collected on the LS-CAT F-Line of the Advanced Photon Source at Argonne National Lab.

The diffraction data was processed and scaled using XDS (version Jun 30, 2023)^46^, the structure was solved by molecular replacement using PHENIX version 1.21^47^ phaser with a model of Ab-RLD generated using ColabFold^48^ as a search model. The generated model was processed to convert pLDDT scores to B-factors using the AFmodel B-factor conversion tool in PHENIX. The final model was built using consecutive rounds of PHENIX real space refine, followed by manual building in Coot v1.1.02 until satisfactory model quality was reached (Table S3). Structure figures were created using UCSF ChimeraX v1.7^49^.

### Size exclusion chromatography for Ab-RLD analysis

Size exclusion chromatography of Ab-RLD was carried out on a Superdex S75 Increase 10/300 GL equilibrated with 20 mM Tris pH 8.0 and 150 mM NaCl. The column void volume was determined using Blue dextran, and calibration elution profiles were generated using the MilliporeSigma low molecular weight protein standards (MWGF70).

### Sulfurtransferase activity assays

Thiosulfate:Cyanide sulfur transfer assays were performed as previously reported^31^. Reactions were carried out in 384 microwell plates using a Tecan H1 Synergy plate reader. The assay buffer contained 20 mM Tris pH 8.0, 150 mM NaCl, and either 50 mM KCN or 5 mM sodium thiosulfate while the respective donor/acceptor concentration was varied from 0 to 600 μM for sodium thiosulfate and 0 to 50 mM for KCN. The reaction was carried out at 25°C, initiated by the addition of Ab-RLD, and stopped after 3 min by adding 1 volume of 20 % (v/v) formaldehyde. The solution was further diluted with 3 volumes of distilled water before adding 1 volume of Goldstein’s reagent^50^ for colorimetric detection of thiocyanate at 460 nm.

Thiosulfate:Glutathione sulfurtransferase assays were performed as previously reported^31^. The assay buffer contained 20 mM Tris pH 8.0, 150 mM NaCl, 0.5 mM lead acetate and 40 mM thiosulfate or 40 mM glutathione (GSH) while varying the respective donor/acceptor concentration from 0 to 10 mM for sodium thiosulfate and 0 to 30 mM for GSH. The reaction was initiated by the addition of 3 μM Ab-RLD, and the formation of lead sulfide was measured by the increase of absorbance at 390 nm in a continuous assay format for 15 min at 25°C using a Tecan H1 Synergy plate reader using 96-well microplates.

### Thermal ramps and stability assays

Thermal ramp assays were performed with an Unchained Labs UNCLE instrument, using the internal fluorescence setup, applying a thermal ramp from 20°C to 95°C at 0.5°C/min. Samples contained 0.5 mg/mL Ab-RLD in 20 mM Tris pH 8.0 and 150 mM NaCl, and either 1 mM sodium thiosulfate, 1 mM GSH, 2 mM sodium sulfate, or 2 mM sodium phosphate. Data analysis was carried out using the Unchained Labs UNCLE analysis tool. Thermal denaturation curves were produced by recording the ratio of fluorescence at 350 nm and 330 nm (350 nm / 330 nm) and melting temperature (T_m_) was determined by taking the first derivative of the denaturation curve.

### Native mass spectrometry analysis

Native mass spectrometric analysis was carried out at the UM Biological Mass Spectrometry Facility. Full length His_6_-tagged Ab-RLD – without TEV cleavage – was used for native mass spectrometry, as we observed slow proteolysis of the N-terminal extension of Ab-RLD after TEV cleavage. Ab-RLD sample concentration was 60 μM. When thiosulfate and/or cyanide were added, 180 μM thiosulfate and 10 mM cyanide were used and samples incubated at room temperature for 2 h. Subsequently, samples were desalted into a 5 mM ammonium acetate pH 7.5 buffer, using pre-equilibrated 50 μL Zeba Spin Desalting Columns (Thermo Fisher Scientific). The eluate was diluted 10-fold and acidified with formic acid (0.1% (v/v) final concentration). Using direct injection, samples were applied to a Thermo Fisher Scientific Q Exactive UHMR Orbitrap Mass Spectrometer. Data was collected using the following general tune settings: 500-6,000 m/z filtering on the quadrupoles, resolution of 100,000 at m/z = 400, averaged over 40 microscans. The ionization chamber was set to 3.8 kV, sweep gas 10, aux gas 5, sheath gas 25, C-trap gas pressure was set to increased value (arbitrary relative units; instrument specific) to detect more charge states. Data was collected across 40 scans (1 scan = 40 microscans). Deconvolution was carried out with Thermo Fisher Scientific BioPharma Finder 5, using the Sliding Window algorithm, filtering for a minimum of 2-3 charge states across 20 intervals.

### Sulfite detection assays

Sulfite detection assays were carried out as previously described^27^. Briefly, sulfite quantification was based on the formation of the colorimetrically detectable *p*-rosaniline sulfonate from *p*-rosaniline. Assays were carried out in 20 mM Tris pH 8.0 and 150 mM NaCl and contained 5 mM thiosulfate or 5 mM thiosulfate and 50 mM KCN. Reactions were started by the addition of Ab-RLD to a final concentration of 50 μM, incubated at 25°C for 5 min, and subsequently quenched using 1 volume of 0.2 M mercury chloride (HgCl). The quenched reaction was centrifuged at 20,000 *g* for 1 min, and the supernatant combined with a 2:1 mixture of 0.04% *p*-rosaniline in 0.72 M HCl and 0.2% (v/v) formaldehyde. After incubation for 5 min, *p*-rosaniline sulfonate absorbance was detected at 570 nm using a Tecan H1 Synergy plate reader.

## Data availability

Structure coordinates for Ab-RLD have been deposited in the Protein Data Bank (PDB) under PDB ID: 8W14.

## Acknowledgements

We thank the staff at the Center for Structural Biology (CSB) at the Life Sciences Institute at UM, especially Dr. Jennifer Meagher, for technical assistance. The CSB is grateful for support from the U-M Life Sciences Institute, the U-M Rogel Cancer Center, the U-M Medical School Endowment for Basic Sciences, and grants from the National Institute of Health. We thank Dr. Carmen Dunbar for technical assistance with native mass spectrometry experiments at the Biological Mass Spectrometry Facility at UM. We thank Dr. Wenjing Wang for access to a Superdex S75 column. We thank Dr. Shero Lao for technical assistance with mass spectrometry data analysis.

## Author contributions

R. B. and T. W. G. conceptualization; R. B. data curation; R. B. and T. W. G. formal analysis; T. W. G. funding acquisition; R. B. investigation; R. B. methodology; T. W. G. and R. B. project administration; T. W. G. supervision; R. B. and T. W. G. visualization; R. B. and T. W. G. writing – original draft; T. W. G. writing – review & editing.

## Funding and additional information

This work was funded by the National Institute of General Medical Sciences (NIGMS) under grant: R35GM133325.

