## Supporting Information for "Structural and biochemical characterization of an encapsulin-associated rhodanese from *Acinetobacter baumannii*"

### Table of Contents

**Figure S1. Additional computational analyses of RLDs** 2
**Figure S2. Biochemical and structural characterization of Ab-RLD** 3
**Figure S3. Thermal ramps of Ab-RLD in the presence of sulfate and phosphate** 4
**Figure S4. Representative native mass spectra of Ab-RLD samples.** 5
**Table S1. Sequence of the gBlock gene fragment of Ab-RLD used in this study** 6
**Table S2. Amino acid sequence of Ab-RLD used in this study** 6
**Table S3. X-ray data collection and refinement statistics for Ab-RLD** 7

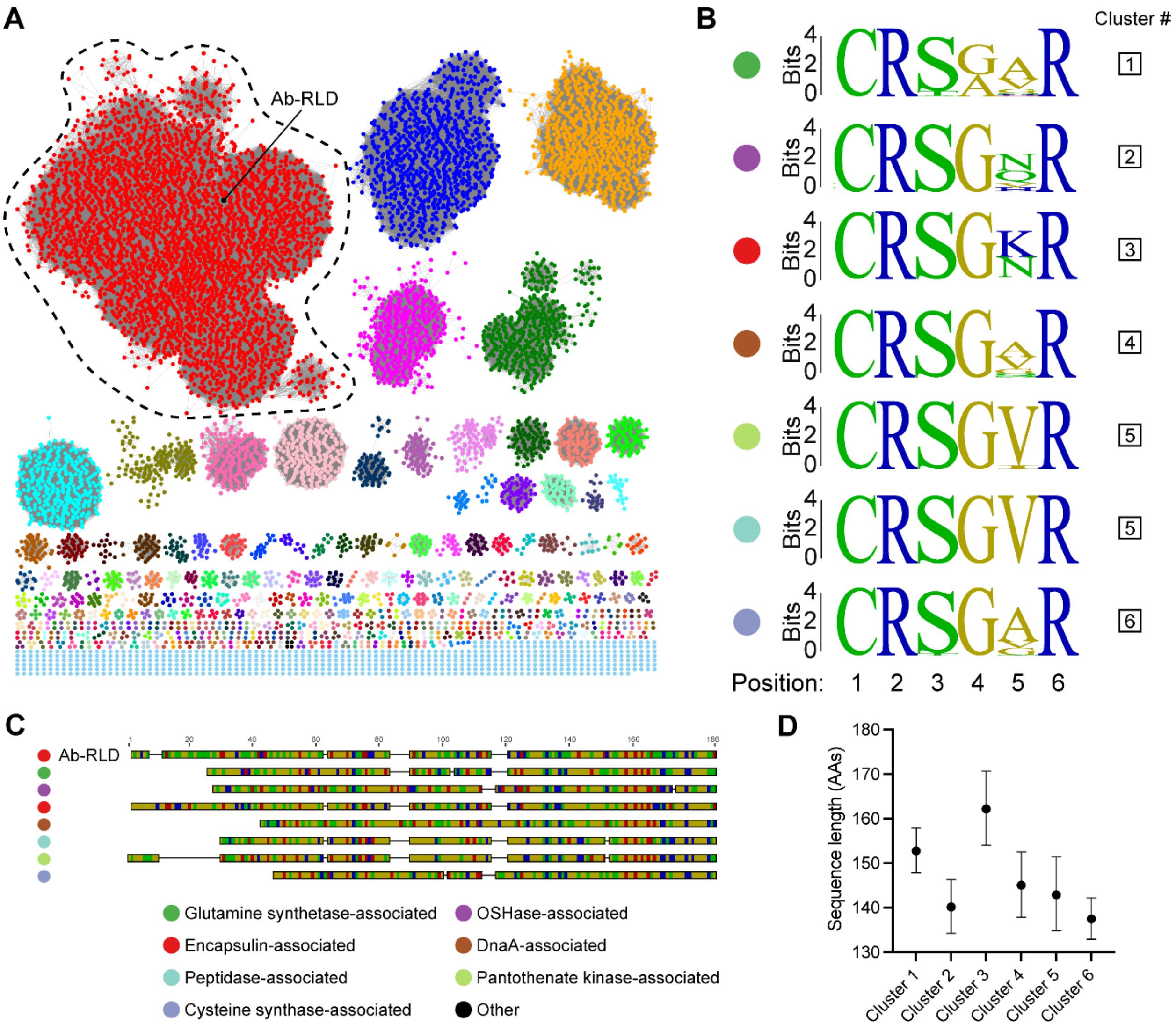


**Figure S1. Additional computational analyses of RLDs.** *A*, initial 10,000-member sequence similarity network (SSN) generated via EFI-EST with Ab-RLD highlighted. The outlined cluster shown in red contains all encapsulin-associated RLDs. *B*, sequence logos of active site loop residues colored by polarity for the six largest clusters identified in the more stringent SSN shown in main Figure 1. *C*, sequence alignment of representative members with sequence length close to the cluster mean from clusters 1-6 compared to Ab-RLD. *D*, mean amino acid sequence length analysis for clusters 1-6 shown in main Figure 1, error bars represent standard deviations from the mean.

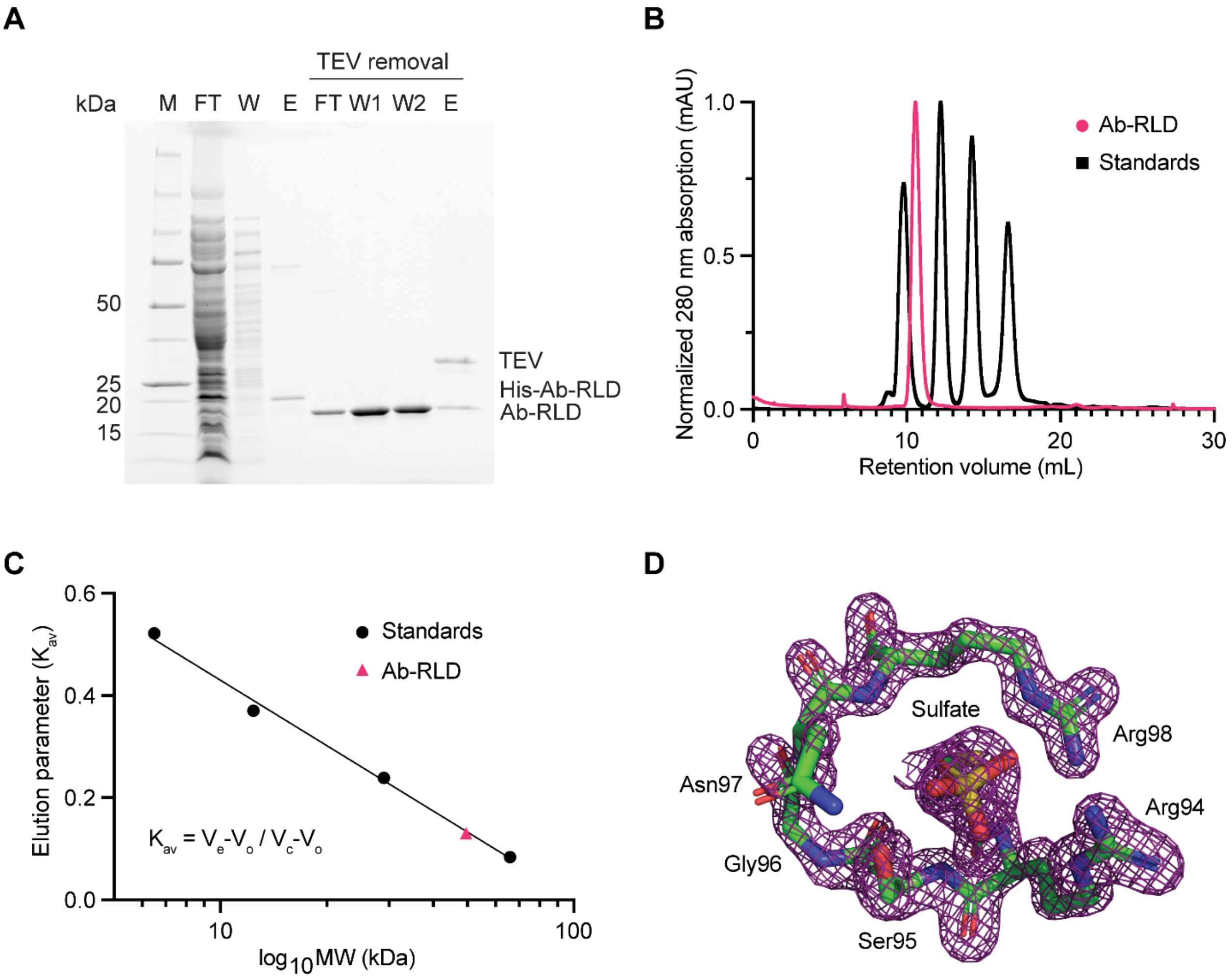


**Figure S2. Biochemical and structural characterization of Ab-RLD.** *A*, SDS-PAGE gel of Ab-RLD purification steps. M: molecular weight marker. FT: Ni-NTA flow through. W: Ni-NTA wash. E: Ni-NTA elution. *B*, size exclusion chromatography analysis of His-tagged Ab-RLD with molecular weight standards (1. bovine serum albumin (BSA) – 66.0 kDa, 2. bovine carbonic anhydrase – 29.0 kDa, 3. horse heart cytochrome c – 12.4 kDa, 4. bovine lung aprotinin – 6.5 kDa) on a Superdex S75 10/300 GL column**.** *C*, size determination of Ab-RLD (ca. 48 kDa) which is most consistent with a dimer (monomer molecular weight: 20.9 kDa). $V_e$: elution volume. $V_o$: void volume. $V_c$: total column volume. *D*, 2Fo-Fc density of sulfate omit map contoured at 2σ showing sulfate modeled into the density in context of the active site loop.

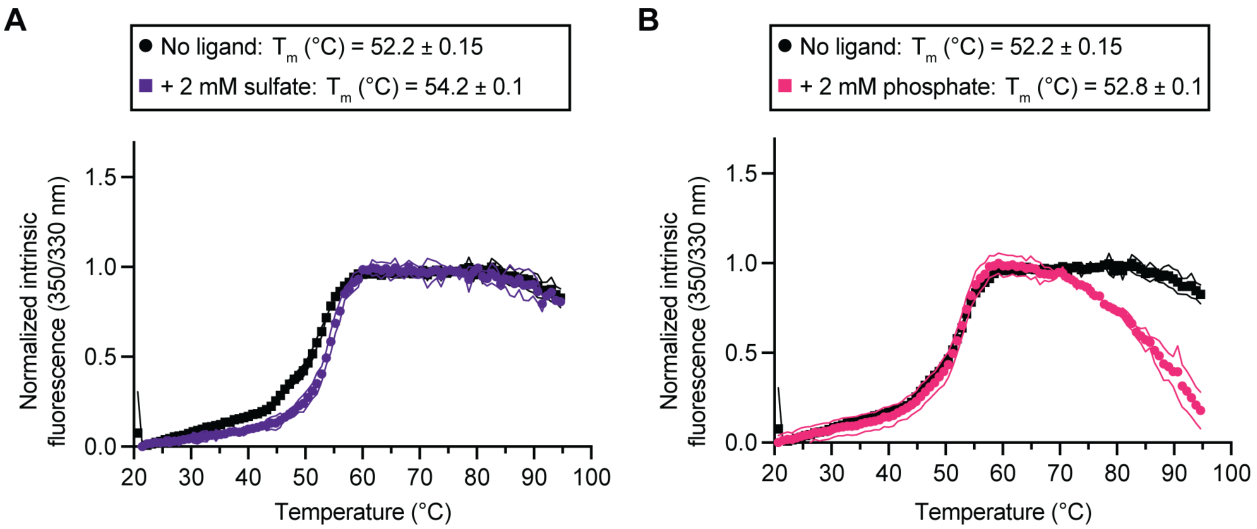


**Figure S3. Thermal ramps of Ab-RLD in the presence of sulfate and phosphate.** *A*, melting curves of Ab-RLD in the presence of sulfate using 350/330 nm intrinsic fluorescence (IF). Each data point represents a normalized mean of 350/330 nm IF. Envelopes highlight standard deviation from the mean. *B*, melting curves of Ab-RLD in the presence of phosphate using 350/330 nm intrinsic fluorescence (IF).

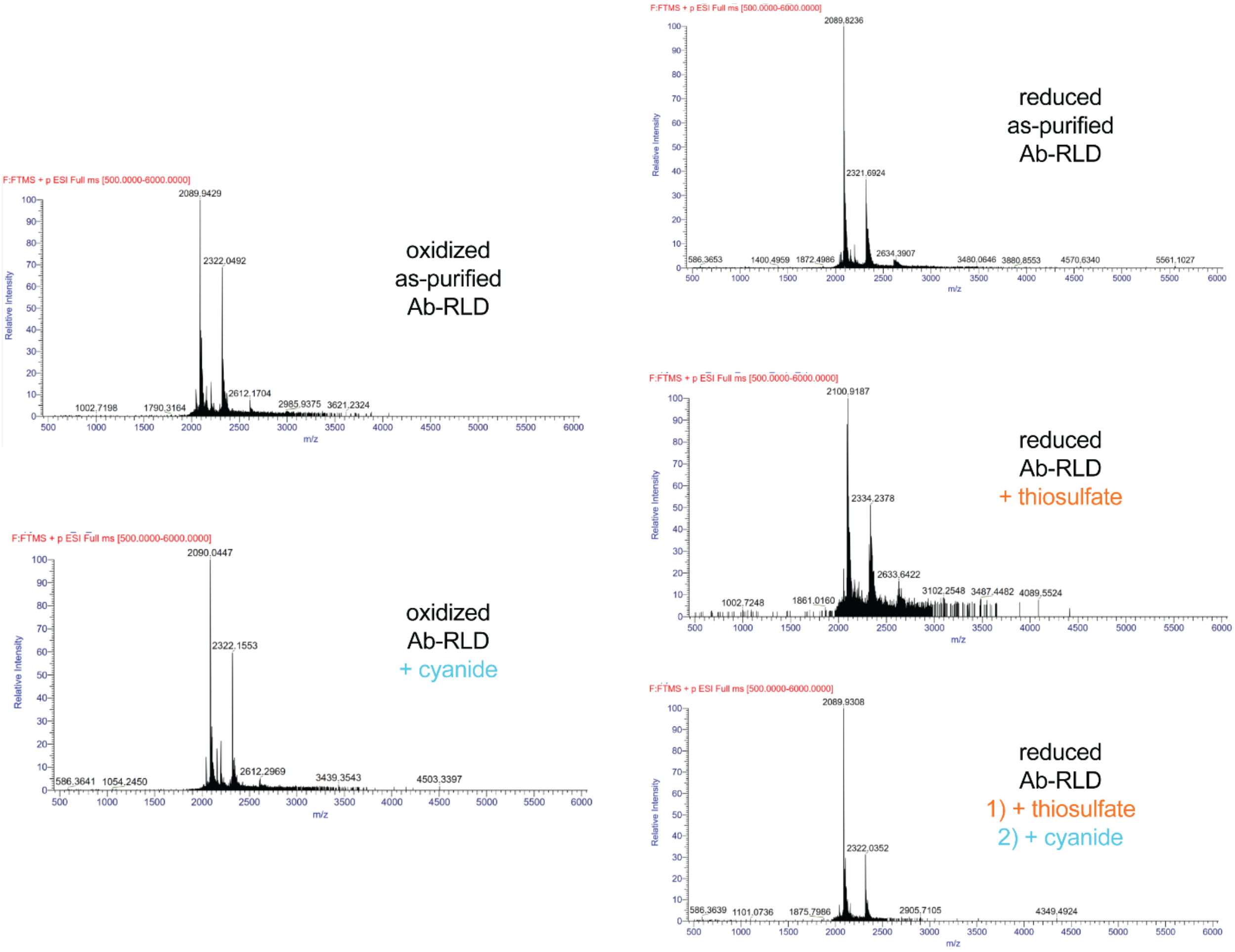


**Figure S4. Representative native mass spectra of Ab-RLD samples.** The respective purification condition (oxidized vs reduced) and donor/acceptor additions are shown.

**Table S1. Sequence of the gBlock gene fragment of Ab-RLD used in this study.**

| Gene | Sequence (Gibson overhangs) |
| --- | --- |
| $His_6$-TEV-GSG-Ab-RLD | AAGTATAAGAAGGAGATATACAATGCATCATCACCACCACCACGA<br>AAATTTGTATTTTCAGGGCAGTGGTAATGCGAAACTTAATACGGAG<br>ACACCTCAAAGTTTTGTCTCCAACGGGTCGAACACGTCATTCAGTG<br>CTGAGGACATCCTGGCGAAGGCACAGCAGTACGCCCAAGAGCACG<br>AATTAAACTTTTCAGGCTCCCTTTCGCCAGTAGATGCCTGGCAACTT<br>GTGCAACAGGGTGAAGCGGTGCTGGTTGACGTACGTACGAATGAA<br>GAGCGTAAATTTGTGGGCTACGTGCCTGAGTCCATCCACGTTGCGT<br>GGGCTACGGGTACGTCTTTCAACCGTAATCCGCGCTTCTTAAAAGA<br>ATTGGAATCAAAGGTTGGTAAAGACAAGACGATTCTGTTATTATGT<br>CGCTCTGGTAATCGCTCAACACAAGCAGCCGAAGCCGCCTTCAATG<br>CTGGGTTCGAGCATATTTACAACGTCCTTGAAGGATTCGAGGGCGA<br>TCTTAACAACAACAGCAGCGTAACCAAAAGAATGGGTGGCGTATC<br>CATCAACTTCCTTGGCAGCAAGACTAAATTAACCTAGGCTGCTGCC<br>ACC |

**Table S2. Amino acid sequence of Ab-RLD used in this study.**

| Protein | Sequence (N-terminal $His_6$-TEV-GSG-tag) |
| --- | --- |
| $His_6$-TEV-GSG-Ab-RLD | MHHHHHHENLYFQGSGNAKLNTETPQSFVSNGSNTSFSAEDILAKAQQYAQ<br>EHELNFSGSLSPVDAWQLVQQGEAVLVDVRTNEERKFVGYVPESIHVAWA<br>TGTSFNRNPRFLKELESKVGKDKTILLLCRSGNRSTQAAEAAFNAGFEHIYN<br>VLEGFEGDLNEQQQRNQKNGWRIHQLPWQQD |

**Table S3. X-ray data collection and refinement statistics for Ab-RLD (PDB ID: 8W14).**

| Statistic | Value | Statistic | Value |
|---|---|---|---|
| **Wavelength** | 0.97872 Å | **CC(work)** | 0.962 (0.864) |
| **Resolution range** | 39.64 - 1.602 (1.659 - 1.602) | **CC(free)** | 0.952 (0.833) |
| **Space group** | P 43 21 2 | **Number of non-hydrogen atoms** | 2783 |
| **Unit cell** | 69.74 69.74 133.23 90 90 90 | **macromolecules** | 2460 |
| **Total reflections** | 378885 (36794) | **ligands** | 22 |
| **Unique reflections** | 43964 (4279) | **solvent** | 301 |
| **Multiplicity** | 8.6 (8.6) | **Protein residues** | 305 |
| **Completeness (%)** | 99.85 (98.82) | **RMS(bonds)** | 0.007 |
| **Mean I/sigma(I)** | 18.15 (3.22) | **RMS(angles)** | 1.05 |
| **Wilson B-factor** | 22.56 | **Ramachandran favored (%)** | 95.68 |
| **R-merge** | 0.06132 (0.5171) | **Ramachandran allowed (%)** | 4.32 |
| **R-meas** | 0.06529 (0.5501) | **Ramachandran outliers (%)** | 0 |
| **R-pim** | 0.02211 (0.186) | **Rotamer outliers (%)** | 2.7 |
| **CC1/2** | 0.999 (0.907) | **Clashscore** | 5.16 |
| **CC*** | 1 (0.975) | **Average B-factor** | 27.03 |
| **Reflections used in refinement** | 43961 (4279) | **macromolecules** | 25.9 |
| **Reflections used for R-free** | 2198 (214) | **ligands** | 36.05 |
| **R-work** | 0.1890 (0.2595) | **solvent** | 35.63 |
| **R-free** | 0.2079 (0.2736) | | |

Statistics for the highest-resolution shell are shown in parentheses.
